# Drainage-structuring of ancestral variation and a common functional pathway shape limited genomic convergence in natural high- and low-predation guppies

**DOI:** 10.1101/2020.10.14.339333

**Authors:** James R. Whiting, Josephine R. Paris, Mijke J. van der Zee, Paul J. Parsons, Detlef Weigel, Bonnie A. Fraser

**Affiliations:** Department of Biosciences, University of Exeter, Stocker Road, Exeter EX4 4QD, UK; Department of Animal and Plant Sciences, University of Sheffield, Alfred Denny Building, University of Sheffield, Western Bank, Sheffield S10 2TN, UK; Department of Molecular Biology, Max Planck Institute for Developmental Biology, Max-Planck-Ring 5, 72076 Tübingen, Germany

**Keywords:** Convergent evolution, genomic convergence, guppy, population genomics, adaptation

## Abstract

Studies of convergence in wild populations have been instrumental in understanding adaptation by providing strong evidence for natural selection. At the genetic level, we are beginning to appreciate that the re-use of the same genes in adaptation occurs through different mechanisms and can be constrained by underlying trait architectures and demographic characteristics of natural populations. Here, we explore these processes in naturally adapted high- (HP) and low-predation (LP) populations of the Trinidadian guppy, *Poecilia reticulata*. As a model for phenotypic change this system provided some of the earliest evidence of rapid and repeatable evolution in vertebrates; the genetic basis of which has yet to be studied at the whole-genome level. We collected whole-genome sequencing data from ten populations (176 individuals) representing five independent HP-LP river pairs across the three main drainages in Northern Trinidad. We evaluate population structure, uncovering several LP bottlenecks and variable between-river introgression that can lead to constraints on the sharing of adaptive variation between populations. Consequently, we found limited selection on common genes or loci across all drainages. Using a pathway type analysis, however, we find evidence of repeated selection on different genes involved in cadherin signalling. Finally, we found a large repeatedly selected haplotype on chromosome 20 in three rivers from the same drainage. Taken together, despite limited sharing of adaptive variation among rivers, we found evidence of convergent evolution associated with HP-LP environments in pathways across divergent drainages and at a previously unreported candidate haplotype within a drainage.

## INTRODUCTION

The process of adaptation in nature can be thought of as a complex interplay between random happenstance and repeatable processes in independent lineages. The latter of these, often termed convergent or parallel evolution [1,2], has provided a myriad of examples from which general rules and principles of adaptation have been dissected under natural conditions. Empirical evidence accumulated over the last decade has demonstrated that convergent phenotypes are often underwritten by convergent changes at the genetic level across many taxa (reviewed in [3–6]). We are now at the point where we can ask why genetic convergence ranges from common in some systems to non-existent in others.

While, there are many definitions of genetic convergence (or parallelism) [2], here we use it broadly to describe selection acting at any of three levels: on the same mutations (eg. [7–9]); different mutations in the same genes (eg. [10–12]); or different genes in the same functional pathways (eg. [13–16]). Further, variation among lineages may arise through one of three modes: either *de novo* mutations (eg. [17,18]); as shared ancestral variation (eg. [19,20]); or through introgression among lineages (eg. [21–24]). An emerging trend within systems is that adaptation involving multiple traits can involve combinations of these levels and modes of convergence. For example, stickleback adapting to freshwater experience selection on ancestral *eda* haplotypes [25] and *de novo* mutation at the *pitx1* gene [10] to repeatedly evolve freshwater bony armour plate and pelvis phenotypes respectively. Similarly, Pease et al. [26] found all three modes of convergence occurring across a clade of wild tomato accessions: adaptive introgression of alleles associated with immunity to fungal pathogens, selection on an ancestral allele conferring fruit colour, and repeated *de novo* mutation of alleles associated with seasonality and heavy metal tolerance. Further, the same phenotype may arise in response to the same selection through any of the above modes, as observed in glyphosate-resistance amaranths across North America [27]. In this study, the authors found glyphosate-resistance evolved in one location by introgression and selection on a pre-adapted allele, in another by the fixation of a shared ancestral haplotype, and in a third location through selection on multiple, derived haplotypes.

Given the range of genetic convergence observed across empirical studies, the importance of different contingencies have emerged. These include the redundancy in the mapping of genotype to phenotype [28,29], i.e. how many genetic routes exist to replicate phenotypes? Limitations in this map are expected to upwardly bias reuse of the same genes or mutations, but redundancy can allow for the evolution of different genes in shared functional pathways. In addition, population structure dictates the sharing of adaptive variation among lineages by which selection may act on [30]. Finally, two lineages may experience an aspect of their environment in a similar way, but in a multidimensional sense environmental variation may limit genetic convergence through pleiotropic constraint [31]. This may result in the re-use of genes with minimal constraint and minimal effects on other aspects of fitness, as suggested for *MC1R* in pigmentation across vertebrates [32]. Alternatively, similarity of environments within multivariate space can predict genetic convergence [33–35], whereby consistencies in the multidimensional fitness landscape channel adaptation along conserved paths. These latter two limiting factors may also explain why genetic convergence can vary for the same traits in the same species in global comparisons, for example in comparisons of Pacific-derived vs Atlantic-derived freshwater stickleback [35–37].

It is clear then to understand the complexity by which genetically convergent evolution might emerge requires study systems for which we already have abundant research on interactions between phenotype and environment. Here, we make use of a model of phenotypic convergence, the Trinidadian Guppy (*Poecilia reticulata*), to evaluate genetic convergence in the replicated adaptation of low-predation (LP) phenotypes from high-predation (HP) sources. For approaching 50 years, this system has provided valuable insights into phenotypic evolution in natural populations, including some of the first evidence of rapid phenotypic evolution in vertebrates across ecological, rather than evolutionary, timescales [38,39]. The guppy has since become a prominent model of phenotypic evolution in nature, but accompanying genomic work has only recently begun to emerge.

The topography of Northern Trinidad creates rivers punctuated by waterfalls, which restrict the movement of guppy predators upstream but not guppies themselves. This replicated downstream/HP and upstream/LP habitat has produced convergent HP-LP guppy phenotypes; LP guppies produce fewer, larger, offspring per brood [40,41], differ in shoaling behaviour [42,43], swimming performance [44] and predator evasion [45], and exhibit brighter sexual ornamentation [46]. Rearing second generation HP-LP guppies in a laboratory setting with controlled rearing conditions confirms that much of the life history differences have a genetic basis [47], and additional work has further demonstrated heritability for colour [48,49] and behaviour [50]. Alongside studies of natural populations, the convergent and replicated nature of these phenotypes has been established with experimental transplanting of HP populations into previously uninhabited LP environments [38,51–53].

Here, we examine whole-genome sequencing of five replicated HP-LP population pairs across the main drainages of Northern Trinidad. Previous work looking at HP-LP convergence in natural HP-LP guppy populations using reduced representation RAD-sequencing found some evidence of molecular convergence [54]. This study however only included three natural populations pairs and inferences from RAD-sequencing can be limited by reliance on linkage disequilibrium and an inability to pinpoint specific candidate genes. To comprehensively explore genetic convergence in this system we first examine how genetic variation is distributed across Northern Trinidad by quantifying population structure, between-river introgression, and within-river demography. We then compare and contrast selection scans between HP-LP pairs within each river to detect signals of convergent evolution. Finally, we examine a large candidate haplotype to explain the mode and mechanisms by which convergence may have occurred at this genomic region.

## RESULTS

### Population structure, admixture, and demographic history

Prior to assessing genomic convergence, it is important to contextualise neutral processes such as population structure, introgression and past demography in order to establish expectations regarding how potentially adaptive genetic variation is distributed and shared among populations; this informs on the most likely mode by which genetic convergence may occur in this system.

Both principal component analysis (PCA) and fineSTRUCTURE [55] confirmed that each river’s HP-LP pair formed sister populations and structure between pairs (as river units) was well-defined. Expectedly, the strongest population structure separated rivers by drainage, with PC1 (33.5%) separating rivers from the Caroni drainage (Guanapo [GHP,GLP], Aripo [APHP,APLP], Tacarigua [TACHP,TACLP]]) from the Norhern and Oropouche drainages (Madamas [MADHP, MADLP] and Oropouche [OHP, OLP] rivers, respectively) (Figure 1A,B). PC2 (19.8%) separated out Caroni rivers, highlighting population structure within this drainage is stronger than structure between Madamas and Oropouche, despite these rivers being in separate drainages. Similarly, admixture proportions inferred by fineSTRUCTURE were lower on average between rivers within the Caroni drainage than between Oropouche and Madamas (Figure 1C). This structure within Caroni occurs despite longer branch lengths in Madamas and Oropouche (Figure 1B) suggesting that the ancestral split between Caroni rivers and Northern/Oropouche rivers is more ancient than the splits between Caroni rivers themselves (Figure 1C). We also detected signatures of introgression into APHP from all Madamas and Oropouche populations, categorised as elevated haplotype donor proportions of individuals from these rivers into APHP recipients (Figure 1C).

**Figure 1:**
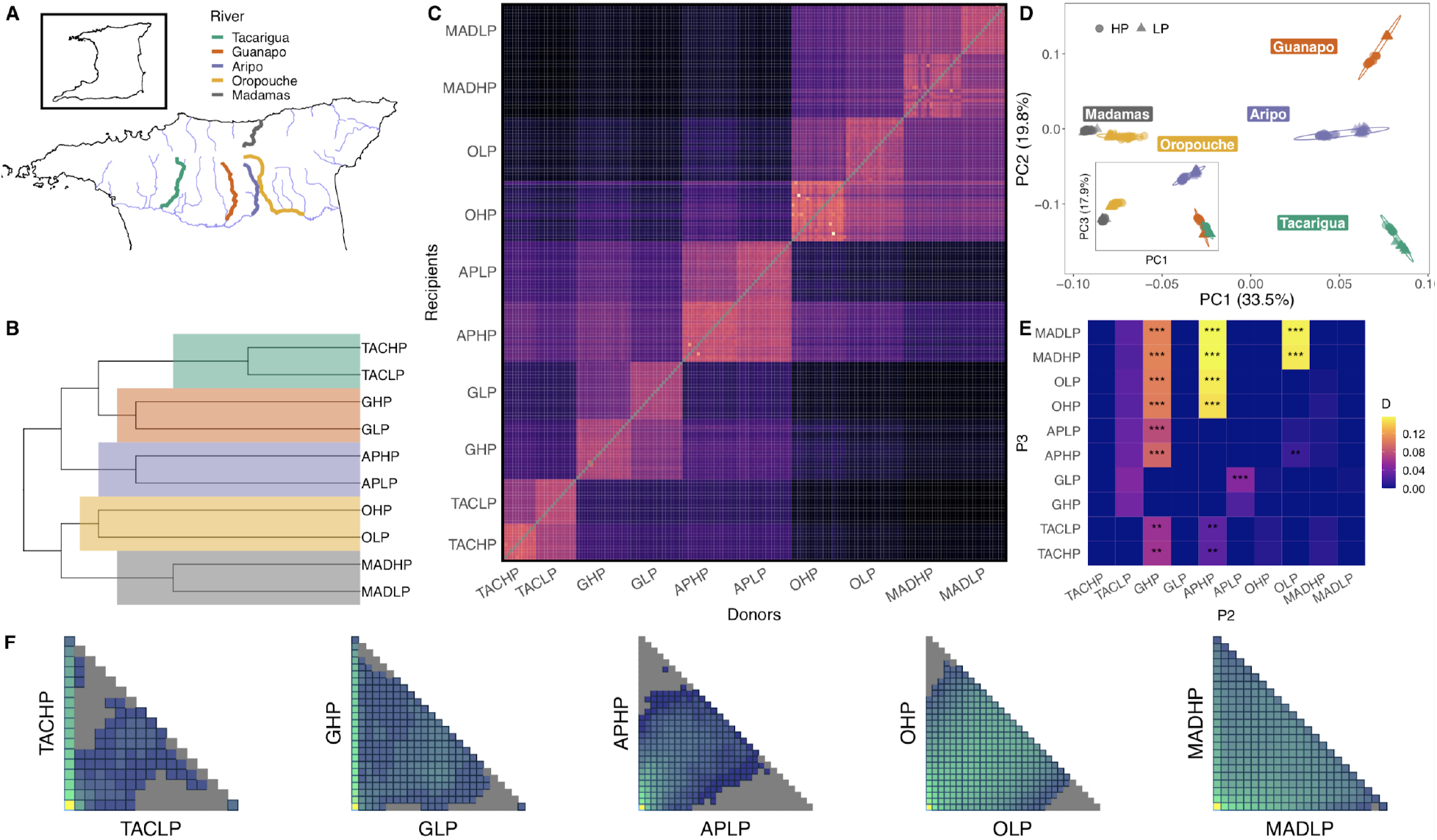
Sampling sites, population structure and admixture across the five rivers. Map (A) highlights sampled rivers from the three drainages in Northern Trinidad. Other major rivers are illustrated in light blue. Sampling rivers plotted alongside topography is available in Figure S1. Phylogenetic tree (B) based on chromosome painting and fineSTRUCTURE analysis, with river nodes highlighted. Heatmap (C) illustrates the proportion of painted recipient haplotypes based on donor haplotypes, used by fineSTRUCTURE to infer coancestry. PCA (D) with populations coloured according to river and shaped according to predation regime. PC3 and PC4 are presented in the insert. Heatmap (E) highlights introgression as pairwise, genome-wide D values between P2 and P3 populations. Populations within rivers were defined as P1 and P2, so all P2 and P3 pairings are between rivers. Two-dimensional site frequency spectra (2dsfs) for each river pair (F) highlighting the sharing of variants between LP and HP populations within each river. LP populations are on the x-axis and HP populations are on the y-axis. In each sfs, the frequency of sites in each population is illustrated from 0 to N, where N is the number of individuals in each population.

To quantify genome-wide introgression, we calculated D-statistics for trios with *Dsuite* (version 0.3; [56]). These were used primarily to infer introgression between populations in different rivers, therefore we focussed on trios where P1 and P2 were HP-LP pairs from the same river. We recovered signals of significant genome-wide introgression in the form of D-statistics (elevated sharing of derived sites between rather than within rivers) between APHP and both populations in the Madamas and Oropouche rivers (D_MADHP_ = 0.156, *p*_MADHP_ = 0; D_MADLP_ = 0.156, *p*_MADLP_ = 0; D_OHP_ = 0.147, *p*_OHP_ = 0; D_OLP_ = 0.152, *p*_OLP_ = 0) (Figure 1E). Comparing donor and recipient haplotype proportions (Figure 1C) suggests that introgression has occurred at a greater rate into, rather than out of, APHP. Further introgression was observed between Madamas populations and OLP (D_MADHP_ = 0.154, *p*_MADHP_ = 0; D_MADLP_ = 0.152, *p*_MADLP_ = 0). Haplotype proportions suggest directionality is greater into OLP. We also observed weak introgression between both Tacarigua populations and APHP (D_TACHP_ = 0.030, *p*_TACHP_ = 6.12e-3; D_TACLP_ = 0.030, *p*_MADLP_ = 6.94e-3), and more likely into APHP. Interestingly, we also detected significant introgression between GLP and APLP (D = 0.05, *p* = 1.17e-7). Evidence of an extreme bottleneck in GLP (see below) is the probable driver of elevated D statistics in comparisons with GHP (Figure 1E), through a loss of within-river shared sites and excessive drift of *de novo* variants in GLP. To detect introgression in spite of this between APLP and GLP suggests the introgression we detected may be conservative.

To assess within river population demography (i.e. LP population bottlenecks, HP-LP migration), we performed demographic modelling based on two-dimensional site frequency spectra (2dSFS) using *fastsimcoal2* [57](Figure 1F; Table 1). All demographic models performed better with the addition of migration, which in every case was higher downstream from LP to HP. For three rivers (Aripo, Madamas, Tacarigua) a historic LP bottleneck was detected alongside stable current population size. Guanapo was better supported by a model with no HP population growth, and Oropouche by a model that suggested HP population growth. The particularly high estimates of N_e_ in APHP agree with the above analyses of introgression into this population.

**Table 1:** Demographic parameters of each river, inferred by *fastsimcoal2*. Values in brackets represent confidence intervals of 95% after bootstrapping 100 SFS.

| Drainage | River | Mean HP-LP $F_{ST}$ | HP $N_e$ | LP $N_e$ | HP > LP Migration | LP > HP Migration | Model |
| --- | --- | --- | --- | --- | --- | --- | --- |
| Caroni | Tacarigua | 0.243 | 3,549 (2864-4493) | 474 (347-612) | 4.78e-5 | 1.55e-4 | LP Bottleneck |
| Caroni | Guanapo | 0.269 | 19,698 (19592-19698) | 155 (135-183) | 3.88e-6 | 8.34e-4 | No population changes |
| Caroni | Aripo | 0.072 | 43,354 (33767-59439) | 6,122 (4588-9008) | 2.10e-4 | 3.27e-4 | LP Bottleneck |
| Oropouche | Oropouche | 0.087 | 4,514 (4148-4910) | 3,740 (3825-4411) | 2.43e-4 | 1.21e-3 | HP population growth |
| Northern | Madamas | 0.171 | 1,121 (922-1355) | 2,480 (2337-2743) | 8.68e-5 | 4.88e-4 | LP Bottleneck |

Altogether, these analyses illustrate how genetic variation is segregated across the five rivers in our dataset. Primarily, ancestral variation is dictated by geography, with populations defined within rivers, then within drainages. Particularly strong population structuring is observed in the Caroni drainage (Tacarigua, Guanapo and Aripo rivers), with limited evidence of introgression having occurred among these rivers within drainage. In contrast, we detect significant introgression across drainages, between the Madamas, Oropouche, and Aripo rivers, demonstrating that these rivers are likely to share a greater proportion of genetic variation than their drainage-structured phylogenetic history would otherwise suggest. These results are consistent with the potential for introgression to occur between rivers through flooding, as the mountainous regions between Tacarigua, Guanapo and Aripo rivers in the Western Caroni drainage are higher and therefore would experience less connectivity during flooding events (Figure S1).

Within rivers, we see evidence of population bottlenecks in LP populations, potentially limiting the amount of available adaptive variation. This is particularly apparent in Tacarigua and Guanapo, within which LP populations have particularly low N_e_ estimates, an excess of monomorphic sites that are polymorphic in the HP founder, and only limited unique polymorphic sites (Figure 1E). In other words, the variation within these LP populations is a subset of that found in the corresponding HP. For all river pairs, our demographic modelling agrees that migration upstream from HP to LP is weaker than LP to HP, compounding the potential for limited variation upstream. Some HP-LP populations are better connected by migration however, such as OHP and OLP, where many polymorphic sites are readily shared between upstream and downstream.

### Candidate HP-LP regions and assessing convergence

To evaluate regions associated with HP-LP adaptation, we scanned the genome using several approaches: XtX [58], a Bayesian analogue of F_ST_ that includes a simulated distribution under neutrality; AFD, absolute allele frequency difference, which scales linearly from 0-1 between undifferentiated and fully differentiated [59]; and XP-EHH (extended haplotype homozygosity) [60], which compares homozygosity between phased haplotypes between populations. To identify selected regions, we calculated each measure in non-overlapping 10kb windows within each river between HP and LP sites. Putatively selected windows were identified if they were detected as outliers by at least two approaches (see methods for outlier criteria for individual tests; Figure S2). Using an intersect of all three may be over-conservative, for example, we would miss instances where divergent selection within a river fixes alternate haplotypes and so both HP and LP population have similarly low heterozygosity (i.e. no XP-EHH outlier but an outlier in XtX and AFD).

Comparing the intersecting list of candidates of XtX, AFD and XP-EHH within each river revealed little overlap among rivers, with only a single 10kb window overlapping in more than two rivers (Figure 2A; for genome-wide plots see Figure S3-5). We then scanned the genome further with 100kb sliding windows (50kb increments) to assess potential clustering of outlier windows in larger regions, but this approach similarly revealed little overlap among rivers. We then explored whether outlier regions (10kb windows overlapping in >1 selection scans) were enriched for genes in common biological pathways between rivers using one-to-one zebrafish orthologues, which may suggest repeated pathway modification, albeit through different genes. Using the outlier regions defined above, no pathway was significantly overrepresented in any river. We did however notice cases in which the same pathways exhibited fold-enrichments >1 in multiple rivers (Figure 2B), albeit non-significant within rivers in each case. We used permutations to explore the likelihood of observing fold-enrichment >1 in our five independently-derived outlier sets. This analysis identified that Cadherin-signalling pathway genes were overrepresented across all five rivers relative to by-chance expectations (*p* = 0.013) (Figure 2B). In total, 20 genes from the Cadherin-signalling pathway were recovered from all five river outlier sets, with some overlap between them (Table S1). This analysis may be over-conservative, due to analysing only guppy genes with one-to-one zebrafish orthologues. Other genes associated with cadherin-signalling were detected by our selection scans, including *Cadherin-1* and *B-Cadherin* in a differentiated region on chromosome 15 (^~^ 5 Mb) in Oropouche and Tacarigua, but these genes exhibited a many-to-many orthology with zebrafish genes so were omitted. We also examined pathways with fold-enrichment >1 in any four rivers, but these were not significant (*p* > 0.05) according to permutation tests (Figure 2B).

**Figure 2:**
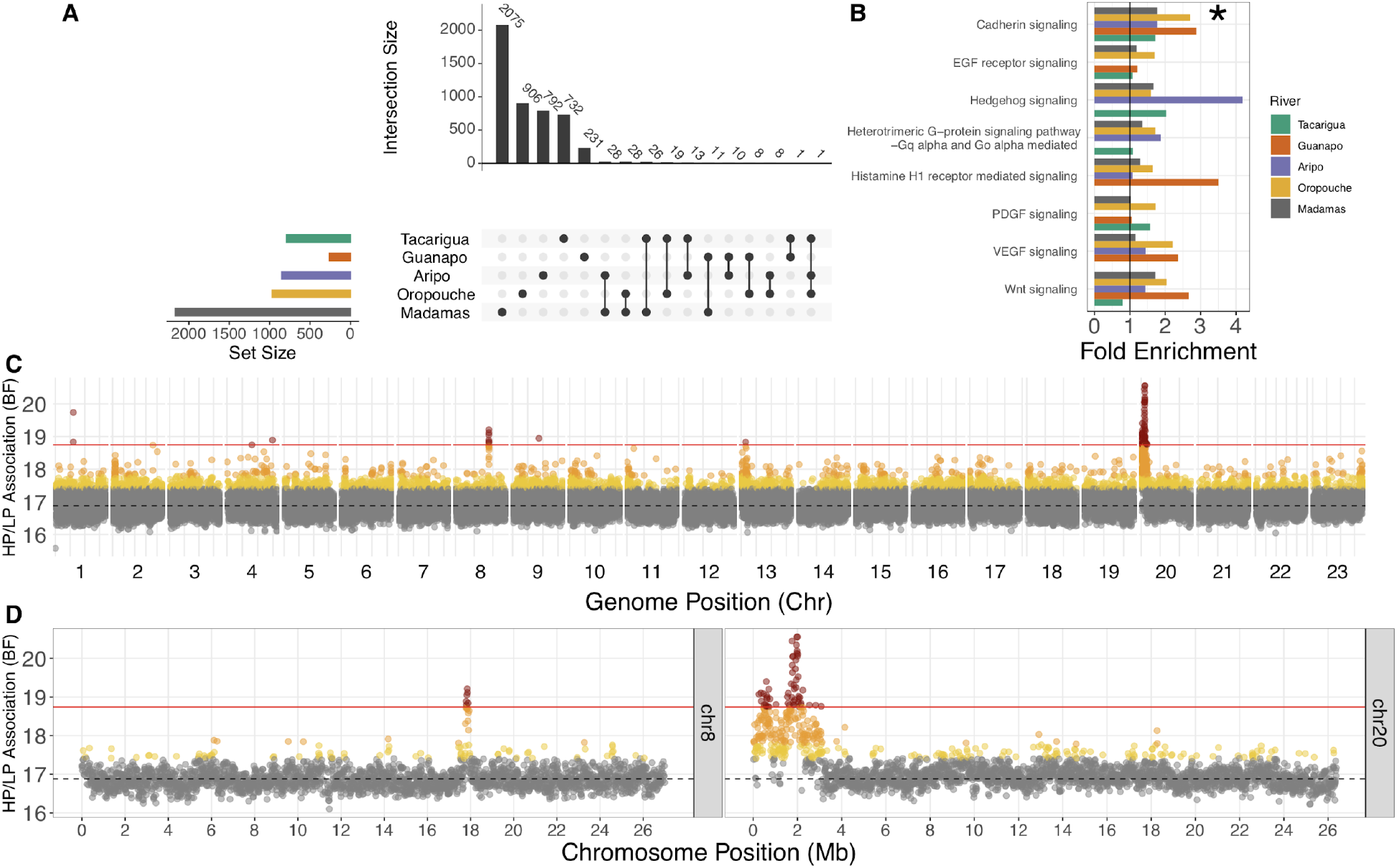
Selection scan results and evidence of convergence. Upset plot (**A**) of overlap among outlier sets (evidence from two of AFD, XtX, XP-EHH) highlighting no overlap beyond sets of three rivers. Fold enrichment of pathways (**B**) associated with zebrafish orthologs of genes in outlier regions within each river calculated from the Panther DB. Cadherin-signalling pathway was the only pathway with fold-enrichment >1 in all rivers. Other presented pathways had fold-enrichment >1 in four rivers. Genome-wide BayPass scan results (**C**) scanning 10kb windows for association with HP-LP classification. Points are coloured according to quantiles: 95% = yellow, 99% = orange, 99.9% = red. Dashed line represents the median BF, and the solid red line denotes 99.9% quantile cut-off. Peaks on chromosomes 8 and 20 are also highlighted (**D**).

We next associated allele frequency changes with HP-LP status using BayPass’ auxiliary covariate model. This latter approach has the advantage of using all populations together in a single analysis, whilst controlling for genetic covariance. Scans for regions associated with HP-LP classification identified two major clusters of associated 10kb windows on chromosomes 8 and 20 (Figure 2C and 2D). In total, we highlighted 70 10kb windows corresponding to 24 annotated genes (and a number of novel, uncharacterised genes) (Table S2). Intersecting these windows with within-river candidate regions highlighted that most HP-LP associated candidates reflected within-river selection scan outliers in one to three rivers (Table S3). Selection scans may overlook some of our association outlier windows because differentiation at these loci may be moderate, but rather we are detecting consistent allele frequency changes in the same direction between HP-LP comparisons. Many of the associated windows mapped to a previously unplaced scaffold in the genome (000094F), but we were able to place this at the start of chromosome 20 along with some local rearrangements (Figure S6) using previously published HiC data [61]. From here on and in Figure 2C-D, we refer to this new arrangement for chromosome 20 and scaffold 000094F as chromosome 20.

The clusters on chromosomes 8 and 20 exhibited multiple 10kb windows above the 99.9% quantile of window-averaged BF scores, suggesting larger regions associated with HP-LP adaptation in multiple rivers (Figure 2D). In particular, the region at the start of chromosome 20 spanned several megabases with two distinct peaks. The larger of these peaks reflected the strongest region of differentiation in the Aripo river (Figure S7) (which was minimally differentiated genome-wide, Figure S5). We explored this candidate region further to evaluate: which rivers showed evidence of HP-LP differentiation within these regions; by what mode of convergence these regions had evolved under; and their gene content and probable candidates for HP-LP phenotypes.

### Candidate Region on Chromosome 20

Visualisation of genotypes (Figure 3A) illustrated extended haplotype structures that were consistent with haplotypes spanning the entire chromosome 20 candidate region (Figure 2D). Interestingly, two of the three Caroni LP populations (GLP and TACLP) were fixed or nearly fixed for homozygous ALT haplotypes across the region (Figure 3A). We will refer to this entire region (^~^0-2.5 Mb) as the ‘CL haplotype’ (Caroni LP haplotype). The other haplotype we will refer to as the ‘REF haplotype’ due to its closer similarity to the reference genome. Moreover, a subset of the candidate region was nearly fixed in APLP (between black lines, between ^~^1.53 - 2.13 Mb, referred to as the CL-AP region Figure 3A). This CL-AP region corresponds to both the region of highest genome-wide divergence in Aripo (Figure S7), and the largest peak in our HP-LP association analysis (Figure 2D).

**Figure 3:**
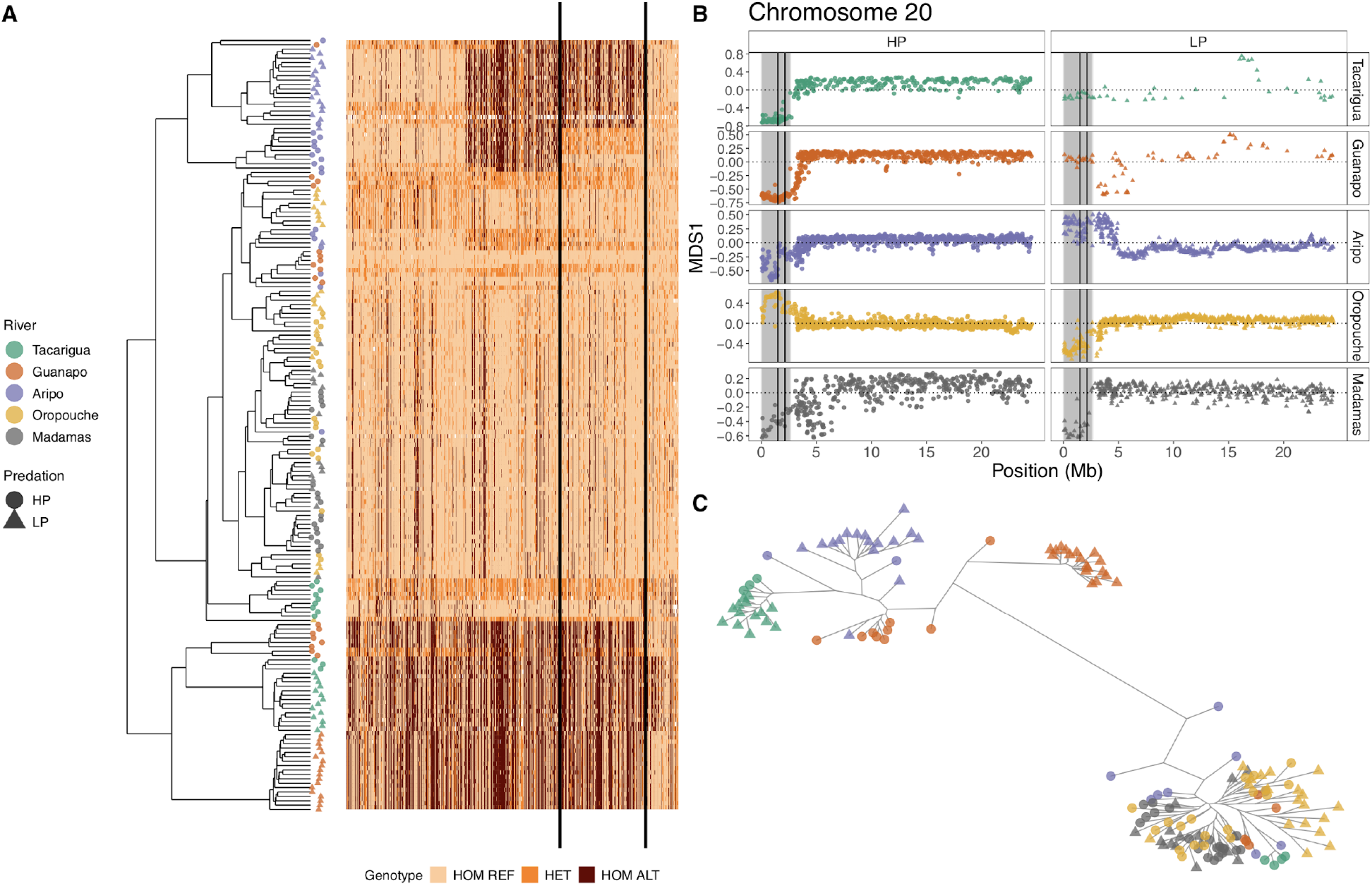
Evidence of divergence along the CL haplotype (ALT alleles, chr20:1-2633448); genotypes for each individual plotted according to hierarchical clustering (**A**). Solid black lines denote the CL-AP region, corresponding to the strongest peak of HP-LP association in the dataset. (**B)** The CL haplotype region (shaded grey) shows evidence of segregated local ancestry in comparison to the rest of chromosome 20, according to MDS scores derived from local PCA analysis. The CL-AP region is again shown between solid black lines. MDS1 = 0 is shown as a dotted line in each panel. A lack of signal on MDS1 for the CL region in Guanapo LP (GLP) and Tacarigua LP (TACLP) reflects that this region is fixed. (**C**) Unrooted maximum-likelihood tree of homozygous individuals (haplogroups) across the CL-AP region, highlighting a major phylogenetic branch separating Caroni LP individuals from both Caroni HP individuals (homozygous for the REF haplotype) and populations from outside the Caroni drainage.

We then assessed whether ancestry of our candidate region (Figure 3B) deviated from the rest of chromosome 20 using a local PCA analysis. This approach is sensitive to inversions, changes in recombination and gene density [62], which may explain why such a large region appears as HP-LP associated. This approach confirmed that the associated region exhibits distinct local ancestry in relation to the rest of the chromosome in all five rivers (Figure 3B), albeit with some idiosyncrasy. For example, MDS scaling along the major axis was broadly similar for GHP and TACHP (LP populations were fixed and therefore showed minimal signal), with the entire region segregated as a single block. In Aripo however, smaller blocks within the region were the major drivers of local ancestry. Linkage analyses confirmed strong linkage across the several megabases spanning the HP-LP associated region in most of the sampled populations (Figure S8).

To understand the mode of convergence at the CL-AP region, we reconstructed the phylogenetic history of the haplotype at this region. To start, we performed PCA analysis over the CL-AP region, and found three clusters along a PC1 axis with large loading (PC1 = 57%, Figure S9), consistent with individuals tending to either be homozygotes for either haplotype or heterozygotes. We used these clusters to define homozygous individuals and explored the phylogenetic history of these homozygote haplotypes following phasing. A maximum-likelihood tree using RAxML-NG (version 0.9.0) illustrated that the CL haplotype at the CL-AP region is phylogenetically distinct from the REF haplotype and separated by a long branch (Figure 3C). This clustering of CL haplotypes and REF haplotypes contradicts the neutral expectation that haplotypes should be predominantly structured within rivers. The clustering of Oropouche and Madamas individuals with Caroni REF haplotypes was surprising, and in stark contrast to the genome-wide phylogeny that clearly separates rivers by drainage (Figure 1B). We compared the branch lengths along this tree among Caroni individuals with the CL haplotype at the CL-AP region with non-Caroni individuals (Oropouche and Madamas) against branch lengths between the same individuals from 50 100kb regions randomly selected from across the genome. This analysis revealed that divergence between the REF and CL haplotype is in line with the modal distance derived from the 50 random trees (Figure S10). This therefore suggests that the CL haplotype may be as old as the split between Caroni and Northern/Oropouche drainage lineages.

These patterns may be consistent with a large inversion, polymorphic in Caroni HP populations but fixed in Caroni LP populations. Further structural variation (SV) or a recombination event between the CL haplotype and the REF haplotype may then have released the CL-AP region from the larger CL haplotype in Aripo only. We therefore explored the potential for inversions and SVs with our aligned read data using smoove [63] (v0.2.5) and Breakdancer [64] (v1.4.5). We did not find evidence for an inversion or an alternative SV spanning either the full CL haplotype (in Guanapo or Tacarigua) or around the diverged CL-AP region in Aripo. Interestingly however, this analysis of SVs did highlight that the strongest peak of HP-LP differentiation in the Oropouche river (chromosome 15 at approximately 5Mb, Figure S11), was associated with a detected 1.1 kb deletion within the *B-cadherin* gene, and exhibited high HP-LP F_ST_ (0.66).

It is particularly interesting that given the CL haplotype may be reasonably old, it has been broken down into smaller regions only in the Aripo river, whereas Tacarigua and Guanapo maintain the full haplotype. One unique attribute of Aripo is evidence of introgression from beyond its drainage. We therefore explored introgression signals (D-statistics) along chromosome 20 between a trio of GH (P1), APHP (P2), and OH (P3) for individuals homozygous for the REF haplotype (upper cluster, Figure 3C). We looked only at these individuals to remove effects of haplotype heterozygosity in GH and APHP but not OH. This analysis strongly suggested that the CL-AP region (and much of chromosome 20) in APHP has experienced introgression with populations outside of the Caroni drainage (greater proportion of shared sites between P2 and P3 compared with P1 and P2) (Figure S12).

Within the CL haplotype region there are 56 annotated genes (Table S4), several of which may have important roles in HP-LP phenotypes. Due to the elevated differentiation observed across the haplotype, it is difficult to pinpoint specific candidates. However, the breakdown of the haplotype in Aripo at the CL-AP region, corresponding to our association peak, provides a unique opportunity to narrow down candidate genes given it is the only part of the larger candidate region that is differentiated in all Caroni HP-LP comparisons. Interestingly, analyses of coverage across the CL-AP region uncovered repeatable low coverage in all Caroni LP populations that corresponded with deletions of several kb (viewed in *igv*) in the LP populations (Figure S13). Five of these deletions overlapped with the *plppr5* gene, including a deletion subsuming the final exon of the gene (Figure S13B). This gene also spanned the HP-LP associated windows with the highest association scores (Table S4). The adjacent *plppr4* gene included the individual SNP (000094F_0:556282) with the highest HP-LP association score of all SNPs in the genome (Figure S13A). This particular SNP was observed in the intron between the second and third exons of *plppr4*.

In summary, this region presents a fascinating example of genetic convergence of an ancestrally-inherited large haplotype among rivers in the Caroni drainage. As such, subsequent genetic divergence of HP-LP adaptation is observed in non-Caroni rivers due to the presumed loss of the CL haplotype in the lineage ancestral to non-Caroni rivers. Additionally, Aripo exhibits a unique signature of stronger HP-LP differentiation at a subregion of the haplotype due to a potentially more recent recombination event between the larger, diverged haplotypes themselves. The CL-AP region exhibits a strong, whole-region signature of introgression with rivers beyond the Caroni drainage (namely neighbouring Oropouche), suggesting that this region of the genome has experienced between-river introgression that may have facilitated haplotype breakdown of the larger CL haplotype.

## DISCUSSION

Using a whole genome sequencing approach, we found a strong candidate haplotype for HP-LP convergence within the Caroni drainage of Northern Trinidad (the only drainage where we have multiple rivers sequenced), but we also found molecular convergence at specific loci is limited among rivers from different drainages. Further, we find evidence that convergence at the level of functional pathways among rivers may facilitate phenotypic convergence across all rivers. Our convergent LP candidate region exhibited a strong signal of divergent selection between HP-LP sites on chromosome 20. This region contains a large haplotype fixed or nearly fixed in LP populations in all three Caroni rivers examined, and contained promising candidate genes for LP phenotypes. This haplotype has largely been maintained as a tightly linked unit, with the exception of Aripo, and likely evolved in these LP populations from shared variation common within, and limited to, the Caroni HP sources. Our analysis of population structure, admixture, and demographic histories across Northern Trinidad suggest that the reduced re-use of the same alleles among drainages may be due to strict structuring of genetic variation between some rivers and recurrent bottlenecks during the founding of LP populations from HP sources. Combined, these processes limit shared ancestral genetic variation from which convergent genetic adaptation may occur. This is not true for all rivers however, with some gene flow taking place between rivers outside of the Caroni drainage and LP populations (APLP and OLP in particular) with no evidence of limited genetic variation.

The candidate region on chromosome 20 was a clear outlier, with by far the strongest signal of HP-LP association, and the majority of windows spanning the first 2.5 Mb of chromosome 20 were above the 99.9% BF cutoff. Interestingly however, most of these windows were only outliers in Aripo for within-river selection scans. Further analysis found allele frequency patterns of a large haplotype nearly fixed in all three LP Caroni drainage rivers (Aripo, Guanapo, and Tacarigua). We termed this large haplotype the ‘CL haplotype’, and within this haplotype the ‘CL-AP region’ as the strongest candidate for convergent HP-LP adaptation due to its particularly strong HP-LP association peak and high within-river differentiation in Aripo. The breakdown of the CL haplotype in Aripo, uniquely, may have been facilitated by introgression from beyond the Caroni drainage. This observation stems from ABBA/BABA D-statistics suggesting the CL-AP region in APHP has experienced introgression with OHP (Figure S12). Recent empirical work has also shown the importance of large haplotype regions containing many genes in convergent evolution. In *Littorina* these large divergent haplotypes are maintained by inversions in crab vs wave ecotypes [65,66]. Similarly, sunflower species repeatedly experience selection on large haplotypes [23], most, but not all, of which are underwritten by inversions. This recent empirical evidence suggests a fundamental role of large haplotype blocks in adaptation by bringing together and maintaining clusters of adaptive alleles, although we cannot rule out genetic draft occurring around a single functional locus within the CL haplotype. We did not detect evidence of inversions within this region, but given that we detect deviations in local ancestry in all rivers (Figure 3B) relative to the rest of the chromosome, and the acrocentric nature of guppy chromosomes [67], it is possible that recombination is reduced over the CL haplotype due to proximity to the telomere. This mechanism could maintain this haplotype in the absence of an inversion. We noted however that, whilst the CL haplotype was fixed (at the CL-AP region) in Caroni LP populations, it was polymorphic in all Caroni HP populations. Large haplotypes may bring together beneficial alleles but they can generate constraint if different loci within the haplotype experience contrasting selection. Through breakdown of the haplotype, potentially facilitated by introgression in Aripo, release of genetic background constraints may have allowed for stronger HP-LP differentiation at the CL-AP region uniquely in this river.

Within the CL-AP region we highlighted the *plppr5* and *plppr4* genes as strong candidates for HP-LP adaptation. These genes correspond to the strongest signals of HP-LP association within the CL-AP region, and in particular the *plppr5* allele on the CL haplotype is associated with an exon-subsuming deletion (Figure S13B). There is limited functional evidence for these genes, but evidence suggests a possible role in growth and body size. Transcriptome analysis has shown that *PLPPR4* is among genes upregulated in slow-growth vs fast-growth Jinghai Yellow Chicken chicks [68], and transgenic mice studies have demonstrated phenotypic effects of *Plppr4* on body size and growth phenotypes [69]. In humans, *PLPPR4* expression is limited to the brain, but *PLPPR5* expression occurs more broadly.

Across the CL haplotype region, HP populations were polymorphic, which is likely why we fail to detect this region as within-river outliers in Guanapo and Tacarigua. Further, these rivers have small LP populations and elevated signatures of genome-wide drift. That the CL haplotype region is variable within HP populations suggests that there is selection on the CL haplotype in LP populations but not against it in HP populations, or that downstream-biased migration is strong enough to maintain the CL haplotype in downstream HP populations. At the CL-AP region, we note that haplotypes derived from Oropouche and Madamas cluster with REF haplotypes from Caroni (Figure 3C), despite genome-wide data suggesting the split between Caroni and Northern/Oropouche drainage rivers is the deepest in our data (Figure 1B). Such patterns can arise when diverged haplotypes are introgressed from more ancient lineages, or even different species, as observed in flatfish [70], sunflowers [23], and *Heliconius* butterflies [24]. We found, however, that branch lengths between the CL and REF haplotype clusters were in keeping with internal branch lengths derived elsewhere in the genome (Figure S10), suggesting it is unlikely that the CL haplotype has evolved elsewhere in an unknown lineage before more recently introgressed into the Caroni drainage. This pattern suggests the CL haplotype may have evolved prior to the splitting of Caroni, Northern, and Oropouche drainage lineages, but was subsequently lost due to drift or selection outside of the Caroni drainage.

By using whole-genome data, we were able to explore in fine-detail how genetic variation is structured and distributed across natural guppy populations across Northern Trinidad. Our observations support previous work suggesting downstream-biased migration, strong drainage-based structuring, and variable gene flow among rivers [71,72]. Using within-river demographic analyses we also found downstream-biased migration in all rivers, but with variable rates, and three LP populations experiencing bottlenecks; these bottlenecks likely represent historical founding bottlenecks as opposed to recent crashes [71]. Such demographic processes, if strong enough, can obscure signals of genetic convergence, or even produce false-positives by manipulating the relative efficacy of selection and neutral processes across the genome [73].

An interesting observation from our introgression analyses was strong signals of introgression in the Aripo HP population from the Oropouche and Madamas rivers. Aripo represents the most easterly river within the Caroni drainage, whilst Oropouche/Quare is the most westerly river in the Oropouche drainage (Figure 1A). Thus, admixture between these populations may be possible, and indeed has been suggested elsewhere [52,72,74]; likely facilitated by flooding during the wet season. We also found weaker, but significant, signals of introgression between APLP and GLP which are upstream and adjacent. Introgression between upstream LP populations has also been reported between the Paria and the Marianne rivers in the Northern drainage [75], but is surprising given the more restrictive topography in the Caroni drainage (Figure S1). The Aripo river may be particularly susceptible to contemporary human translocations of guppies from across Trinidad, as it is heavily involved in active research including experimental introductions.

These population structure results have important implications for the likelihood of observing genetic convergence because they identify constraints on the sharing of ancestral genetic variation through LP founding bottlenecks and limitations on adaptive variants being shared among rivers [3]. Such structuring of genetic variation, typical of riverine populations, stands in contrast to other prominent systems with abundant evidence of genomic convergence, such as sticklebacks [76] and atlantic herring [77], where largely panmictic marine populations share much adaptive variation. Standing genetic variation is a major contributor of adaptive variation [78–80], and the sharing of this variation among lineages acts a significant contingency to genetic convergence [81] in varied systems including fish [25,82] and insects [83].

Redundancy in the mapping of phenotype to genotype is expected for highly polygenic traits, whereby many loci may be adapted to modify a phenotype, and in instances where phenotypes are derived from complex functional pathways [28]. Indeed, convergence at the level of pathways has been described for human pygmy phenotypes [15] and hymenopteran caste systems [14]. In our selection scans, we found a greater proportion of genes associated with cadherin-signalling than expected across five replicated datasets, suggesting this pathway may be under selection at different genes in all rivers. Cadherin genes *cadherin-1* and *B-cadherin* have previously been detected as under selection in experimentally transplanted LP populations derived from GHP [54], however these specific genes, whilst also detected here, were omitted from our pathway analysis due to many-to-one orthology with zebrafish genes. Genes in this pathway have important roles in cell-cell adhesion, and are associated with tissue morphogenesis and homeostasis by mediating interfacial tension and orchestrating the mechanical coupling of contact cells [84]. Cadherin signalling pathways interact with the signalling of various growth hormones and are involved in differential growth phenotypes. For example, cadherin signalling genes were differentially expressed between transgenic and wild-type coho salmon with divergent muscle fibre phenotypes that affect growth and the energetic costs of maintenance [85]. Cadherin genes are also expressed during oogenesis in *Drosophila* [86]. Assessing the functional roles of the cadherin signalling genes identified in our study (Table S1) is beyond the scope of this work, but in identifying this pathway across rivers we provide evidence for a potentially shared mechanism by which HP-LP phenotypes may similarly evolve across Northern Trinidad.

It is also likely that contrasting demographic contexts within rivers, particularly migration between HP-LP, can influence the genetic architecture of traits or the regions of the genome where adaptive alleles reside. Theory predicts that with increasing sympatry, if multiple genetic routes to a phenotype exist then traits should be underwritten by simpler genetic architectures with fewer loci, each of larger effect [87,88]; such that favourable combinations of alleles are less likely to be broken down by introgression. Empirical support from cichlid species pairs suggests this is the case for male nuptial colour traits in sympatric vs allopatric pairs [89]. Given demographic histories vary between our HP-LP populations (Table 1), these natural conditions may moderate the selective benefits of different genetic routes to phenotypes.

In conclusion, we have investigated whether convergent HP-LP phenotypes that have evolved repeatedly within rivers across Northern Trinidad are underpinned by convergent genetic changes. We found convergence of genetic pathways not specific genes across drainages. This is in keeping with recent work suggesting a predominant role of shared standing genetic variation in driving convergent changes at the gene-level, a mechanism that is restricted in natural guppy populations by limited between-river gene flow and recurrent founding bottlenecks during LP colonisations. Within our drainage with multiple rivers sampled, we did however find convergent evolution of a large haplotype region nearly fixed in Caroni LP populations that is likely derived from shared ancestral variation between these populations. These results provide a comprehensive, whole-genome perspective of genetic convergence in the Trinidadian guppy, a model for phenotypic convergence.

## METHODS

### Sampling, Sequencing and SNP calling

Individuals were sampled from naturally-occurring downstream HP and upstream LP environments in rivers from each of Northern Trinidad’s three drainages (Figure 1) between 2013 and 2017. Three of these rivers (Aripo, Guanapo, Tacarigua) share a drainage (Caroni), whilst Oropouche (Oropouche) and Madamas (Northern) are found in separate drainages. Numbers of individuals from each population were: TACHP = 12, TACLP = 14, GHP = 19, GLP = 18, APHP = 19, APLP = 19, OHP = 19, OLP = 20, MADHP = 20, MADLP = 16. Samples were stored in 95% ethanol or RNeasy at 4° C prior to DNA extraction. Total genomic DNA was extracted using the Qiagen DNeasy Blood and Tissue kit (QIAGEN; Heidelburg, Germany), following the manufacturer’s guidelines. DNA concentrations ≥ 35ng/μl were normalised to 500ng in 50μl and were prepared as Low Input Transposase Enabled (LITE) DNA libraries at The Earlham Institute, Norwich UK. LITE libraries were sequenced on an Illumina HiSeq4000 with a 150bp paired-end metric and a target insert size of 300bp, and were pooled across several lanes so as to avoid technical bias with a sequencing coverage target of ≥10x per sample. Data from the Guanapo and Oropouche rivers has been previously published as part of Fraser et al. [61].

Paired-end reads were quality-controlled with fastQC (v0.11.7) and trimmed with trim_galore (v0.4.5) before being aligned to the long-read, male guppy genome assembly [61] with bwa mem (v0.7.17). Appropriate read groups were added followed by alignment indexing, deduplication and merging to produce final bams. Merged, deduplicated alignments were recalibrated using a truth-set of variants generated from high-coverage, PCR-free sequencing data from 12 individuals [61]. GVCFs were produced using GATK’s (v4.0.5.1) HaplotypeCaller and consolidated to chromosome/scaffold intervals with GenomicsDBImport prior to genotyping with GenotypeGVCFs.

SNPs were filtered on the basis of QD < 2.0, FS > 60.0, MQ < 40.0, HaplotypeScore > 13.0 and MappingQualityRankSum < −12.5 according to GATK best practices. We retained only biallelic sites with a depth ≥ 5. SNPs were also removed if missing in > 50% of individuals within a population, if they were not present in all ten populations, and had a minor allele frequency < 0.05 (relative to all individuals). This produced a final dataset of 3,033,083 high-quality SNPs.

### Population structure and introgression

Principal Component Analysis was performed over all ten populations using a linkage-pruned (--indep-pairwise 50 5 0.2) set of SNPs (N = 217,954) using plink (v2.00).

Prior to further estimates of population structure, chromosomes and scaffolds were phased individually with beagle (v5.0; [90]), which performs imputation and phasing, and then phased again using shapeit (v2.r904; [91]) making use of phase-informative reads (PIR). This method has been effective elsewhere [92]. Phased chromosomes were re-merged into a single file for analysis with fineSTRUCTURE (v4.0.1; [55]). FineSTRUCTURE first makes use of chromosome painting before assessing admixture based on recombinant haplotype sharing among all individuals. FineSTRUCTURE was run with a uniform recombination map and an inferred *c*-value of 0.344942 with successful convergence.

We also estimated D-statistics for all 120 possible trios of the 10 populations using Dsuite [56]. Here, sequencing data from 10 *P. picta* individuals (5 males, 5 females) was aligned to the male guppy genome using the protocol above to be used as an outgroup. bcftools consensus was used to create a *P. picta* fasta file based on the final SNP variant calls and ancestral alleles were inserted into the *P. reticulata* VCF files using *vcftools fill-aa*. We highlighted introgression candidates as trios with significant D values (Bonferroni-corrected *p*-value < 0.05) between populations from different rivers i.e. we retained trios for which P1 and P2 were assigned to the same river by including the tree structure in Figure 1B to examine the possibility of introgression between rivers. To examine further introgression along chromosomes, we calculated D-values in non-overlapping windows of 100 SNPs.

### Demographic inference

We explicitly modelled the demographic history of each HP-LP population pair using *fastsimcoal2* [57]. Fastsimcoal2 uses a continuous-time sequential Markovian coalescent approximation to estimate demographic parameters from the site frequency spectrum (SFS). As the SFS is sensitive to missing data, a --max-missing filter of 80% was applied to each population vcf containing both monomorphic and polymorphic variants, and to remove LD, the vcf was thinned at an interval of 20kb using *vcftools* [93]. For each HP-LP population pair, we generated a folded two-dimensional frequency spectrum (2dSFS) using the minor allele frequency, which were generated via projections that maximised the number of segregating sites using easySFS (https://github.com/isaacovercast/easySFS).

Five demographic models were used to explore the demographic history of the population pairs, all of which contained a uniform distribution divergence of 1 to 6e7 and log uniform distribution N_E_ of 1 to 50000: Model A) HP-LP split with no population growth; Model B) HP-LP split with no population growth and a post-founding bottleneck in the LP population; Model C) HP-LP split with historical population growth in HP; Model D) HP-LP split with historical population growth in HP and a post-founding bottleneck in the LP population; Model E) HP-LP split with bottlenecks in both LP and HP populations. All five models were also run with the inclusion of a migration matrix between LP and HP with a log uniform distribution of 1e-8 to 1e-2. Fastsimcoal2 was used to estimate the expected joint SFS generated from 100 independent runs, each consisting of 200,000 simulations per estimate (-n), generated by 100 ECM cycles (-L). Model choice was assessed by computing the log likelihood ratio distributions based on simulating 100 expected SFS from the run with the lowest delta (smallest difference between MaxEstLhood and MaxObsLhood) as per Bagley et al. [94]. The most likely model was selected for each population pair and the run with the lowest delta likelihood was used as input for bootstrapping by simulating 100 SFS. We report the median and 95% confidence intervals for N_E_ and probabilities of migration as provided by bootstrapping.

### Scans for selection

We used three approaches to scan the genome for signatures of selection between HP-LP populations. We first estimated XtX (a Bayesian approximation of F_ST_) within each river using Baypass [58], which has the advantage of including a genetic covariance matrix to account for some demographic variation. Genetic covariance matrices were estimated using LD-pruned (plink --indep-pairwise 50 5 0.2) VCFs for each river, and averaged over 10 independent runs. We determined a significance threshold within each river by simulating neutral XtX of a POD sample of 10,000 SNPs with the *simulate.baypass(*) function in R. We then marked 10kb windows as outliers if their mean XtX value exceeded the 0.95 quantile of the neutrally simulated distribution. Secondly, absolute allele frequency differences (AFD) were estimated by taking allele counts from each population (vcftools --counts2) and estimating frequency changes per SNP. To convert per SNP values to 10kb non-overlapping windows we removed invariants within each river, calculated the median AFD, and filtered windows that contained fewer than 6 SNPs. We marked outliers as windows above the upper 0.95 quantile, or with an AFD > 0.5 if this quantile was > 0.5. The linear association with AFD and differentiation, compared to the non-linear equivalent for F_ST_ makes this comparable measure of allele frequency change more interpretable [59]. A cutoff of AFD = 0.5 therefore represents the minimum by which to observe a change in the major allele between HP and LP. Finally, we estimated the extended haplotype homozygosity score XP-EHH between each river with selscan (v1.2.0a; [95]) and normalised in windows of 10kb. We limited this analysis to chromosomes and scaffolds > 1 Mb in size, due to extreme estimates on smaller scaffolds distorting normalisation. Outliers were marked as those with normalised XP-EHH > 2.

Enrichment analyses were performed by extracting one-to-one zebrafish orthologues for guppy genes in outlier windows using Ensembl’s BioMart (release 101; [96]). Orthologues were then assessed for enrichment within rivers by comparing outlier genes against a background set of all genome-wide guppy-zebrafish one-to-one orthologues using PantherDB [97]. Results from all rivers were then compared in a single analysis to assess the above-expected enrichment of functional groups across the entire dataset. We performed random permutations (N=10,000) to draw equivalently sized within-river sets of outliers with weightings based on the number of genes within each functional pathway group within the guppy-zebrafish orthologue background gene set. Based on permuted random outlier sets, we then assessed the by-chance likelihood of observing within-river enrichment >1 for all five or any four rivers for each of the functional groups where this had been observed (groups plotted in Figure 2B).

For associated allele frequency changes with HP-LP classification, we also applied BayPass’s auxillary covariate model, which associates allele frequencies with an environmental covariate whilst accounting for spatial dependency among SNPs with an Ising prior (-isingbeta 1.0). We used a genetic covariance matrix including all 10 populations estimated as above. We split our SNP data into 16 subsets, allocating alternate SNPs to each subset such that all subsets included SNPs from all regions of the genome, and merged outputs as recommended in the manual. We averaged per SNP BayesFactor scores within 10kb and marked as outliers those above the 0.999 quantile. We also explored alternative windowing strategies here, such as marking outlier SNPs above the 0.999 quantile and determining window significance as windows with significantly more outlier SNPs than a binomial (99.9%) expectation, however this made a minimal impact on which windows were deemed outliers.

### Characterisation of structural variants

To assess whether candidate regions may contain structural variants, particularly inversions, we first used local PCA analysis within each population for each chromosome with the R package *lostruct* (v0.0.0.9) [62]. This method explores phylogenetic relationships between individuals using windows of N SNPs along a chromosome, and then uses multidimensional scaling to visualise chromosomal regions that deviate from the chromosomal consensus among individuals. We filtered each population’s chromosome for invariants, prior to running local PCA in windows of 100 SNPs. For each run, we retained the first two eigenvectors (k=2) and computed over the first two PCs (npc=2).

We used two methods to call structural variants from our final bam files: smoove (v0.2.5; [63]) (a framework utilising lumpy [98]) and Breakdancer (v1.4.5; [64]). For both methods we called structural variants across all HP and LP individuals within a river. For smoove variants, we excluded repetitive regions of the genome prior to variant calling, and filtered the subsequent VCF for SVs marked as imprecise; <1kb in size; less read pair support (SU) than the per-river median; without both paired-end and split-read support (PE | SR == 0). Breakdancer was run per river, per chromosome, with results filtered for SVs <1kb in size; less support than per river, per chromosome median support; quality < 99. We calculated F_ST_ using smoove VCFs within each river to explore structural variants that may have diverged within rivers. To validate SVs of interest, we plotted all bam files within a river using samplot [99] and visualised regions in *igv*.

### Phylogenetic relationships of haplotypes

To examine phylogenetic relationships of the CL haplotype, we calculated maximum likelihood trees among homozygote haplogroups using RAxML-NG (v0.9.0; [100]). We limited this analysis to the CL-AP region, which was subsumed within the larger CL haplotype in Guanapo and Tacarigua and included the strongest region of HP-LP association (CL-AP region). Haplogroups were defined on the basis of PC1 clusters (Figure S9). We retained a random haplotype from homozygote individuals from all populations and constructed a tree using the GTR + Gamma model with bootstrap support added (500 trees) with a cut-off of 3%.

To compare against other regions of genome, we randomly sampled 50 regions of the genome of 100kb for the same individuals, repeated the above method for tree calculations, and estimated branch lengths between individuals Caroni drainage individuals from the CL haplogroup and Oropouche/Madamas individuals from the REF haplogroup.

## AUTHOR CONTRIBUTIONS

JRW wrote the manuscript and performed all statistical and genomic analyses with the exception of demographic modelling, which was done by JRP and MJvdZ. JRP, MJvdZ and PP were involved in wet lab work including DNA extraction and library preps. DW procured grants for sampling. BAF conceived and supervised the project, wrote and edited parts of the manuscript, undertook sampling in Trinidad and was responsible for all other funding.

## ACKNOWLEDGEMENTS

Kim Hughes and Mitchel Daniel for *Poecilia picta* specimens. David Reznick for support in sampling logistics. HPC infrastructure support was provided by The University of Exeter’s High-Performance Computing (HPC) facility (ISCA). DNA sequencing was performed by University of Exeter Sequencing Service (ESS). Batch submission scripts for *fastsimcoal2* analyses were kindly provided by Vitor Sousa.

## DATA ACCESSIBILITY (To be updated upon acceptance)

Population genomics data are available on ENA: Study: XXX

Scripts used to analyse data are available from an archived github repository: Zenodo DOI: XXX

