## Supplementary material for "Drainage-structuring of ancestral variation and a common functional pathway shape limited genomic convergence in natural high- and low-predation guppies": Figure S

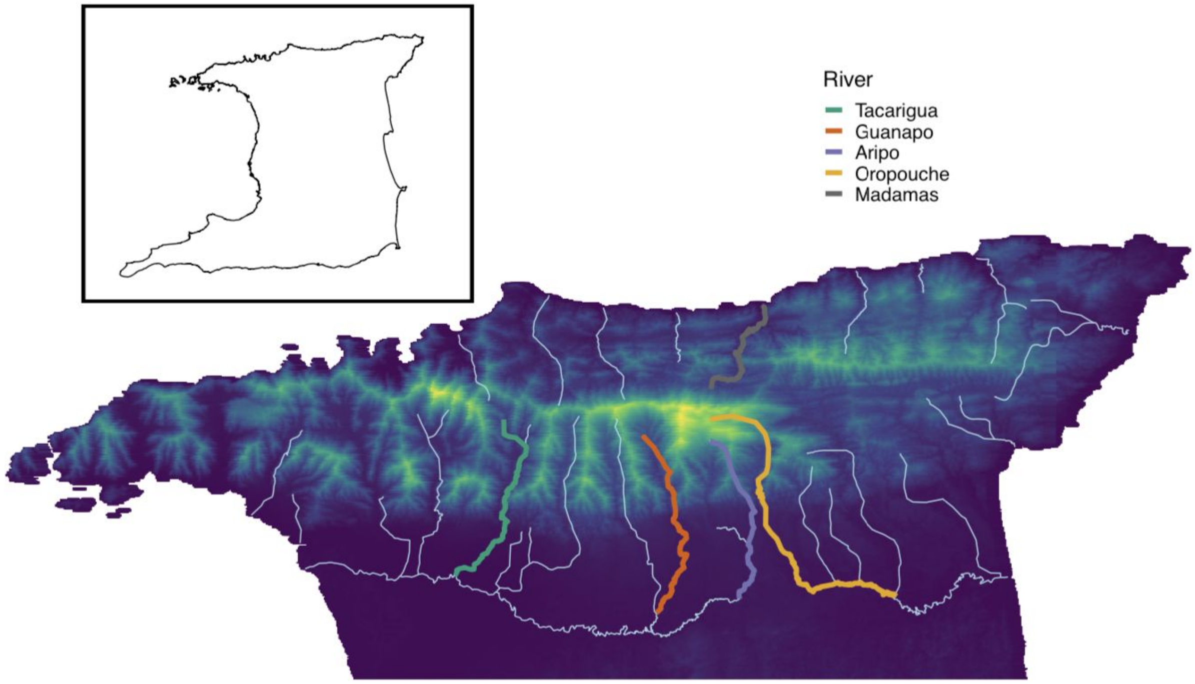


**Figure S1:** Map of sampling rivers in Northern Trinidad alongside topography. Sampling rivers are coloured according to legend and other major rivers are coloured light blue. Topography ranges from low altitude (dark fill) to high altitude (light fill). Upstream regions of rivers in the Western Caroni drainage (Tacarigua, Guanapo and Aripo) are flanked by mountain ranges, likely preventing gene flow occurring between these rivers.

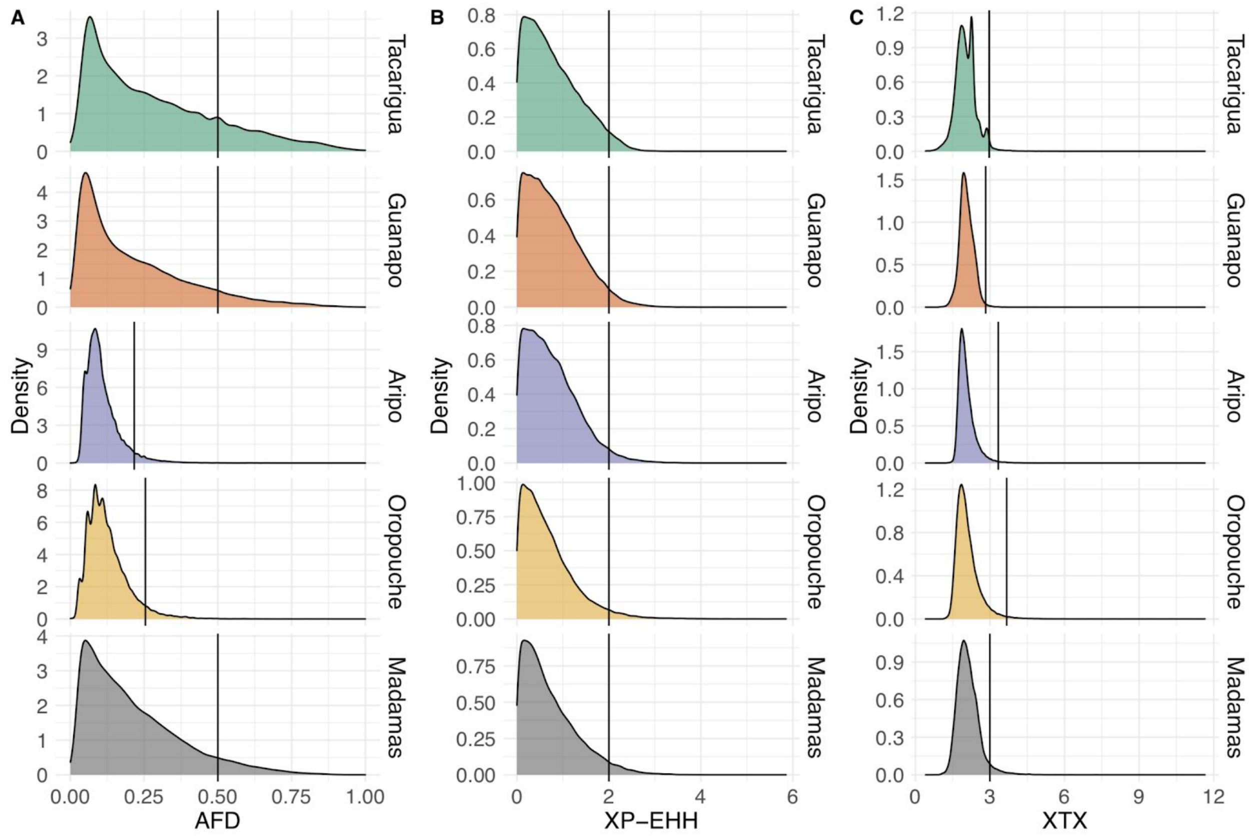


**Figure S2:** Distributions of selection scanning methods within each river and their associated outlier cut-offs for AFD (**A**), XP-EHH (**B**) and XtX (**C**).

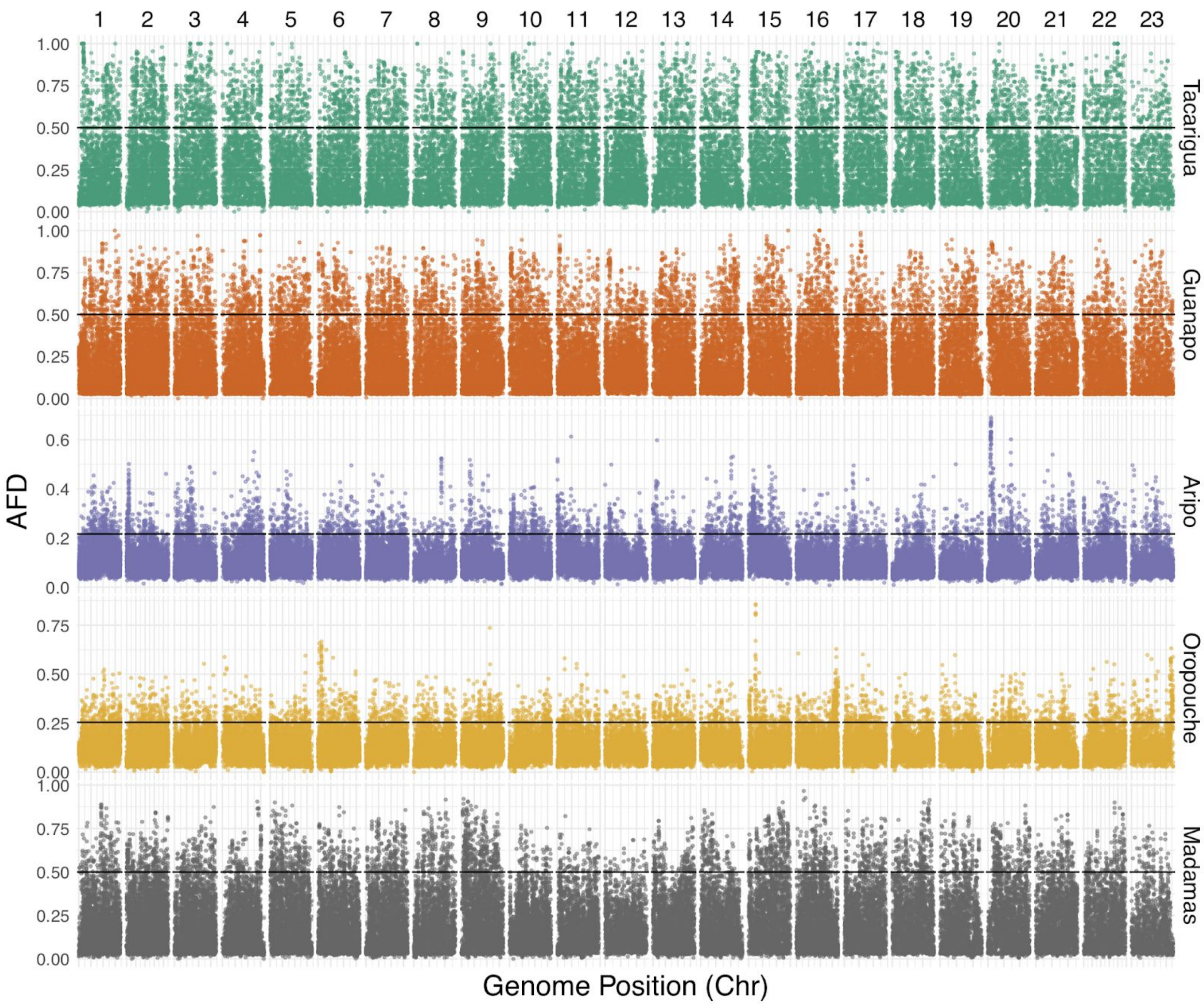


**Figure S3:** Genome-wide AFD results for 10kb windows. Panels represent the 23 chromosomes in the guppy genome. Each row represents the change in the absolute allele frequency for 10kb windows between HP and LP populations in a different river. Chr20 has been updated to include the unplaced scaffold 000094F. The horizontal line in each row denotes river-specific outlier cutoffs.

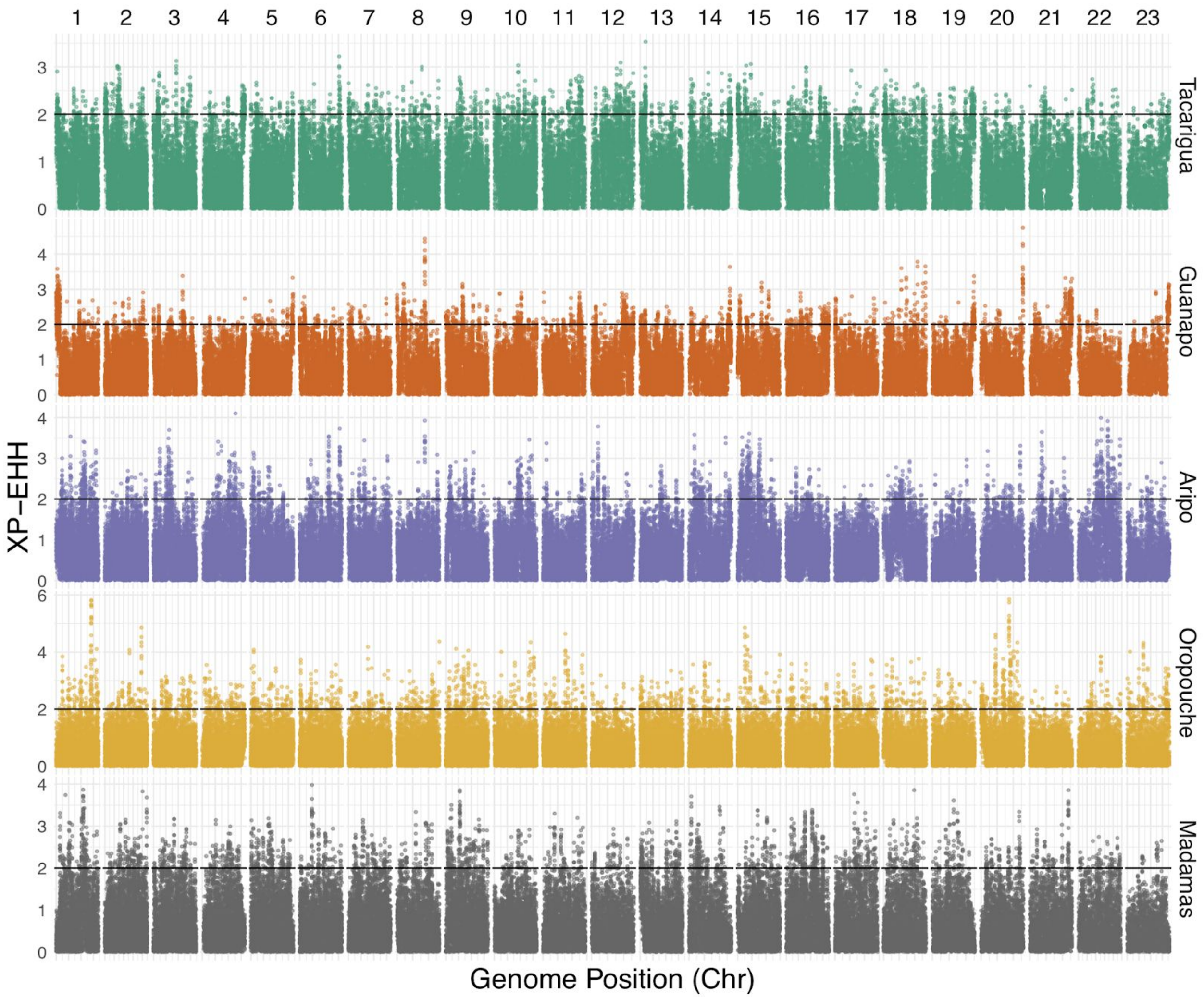


**Figure S4:** Genome-wide XP-EHH results for 10kb windows. Panels represent the 23 chromosomes in the guppy genome. Each row represents the normalised score for XP-EHH, which compares extended haplotype homozygosity between HP and LP populations within rivers (absolute-transformed). Chr20 has been updated to include the unplaced scaffold 000094F. The horizontal line in each row denotes the outlier cutoff = 2, analogous to a Z-score > 2 reflecting approximately $p$ = 0.05 following normalisation.

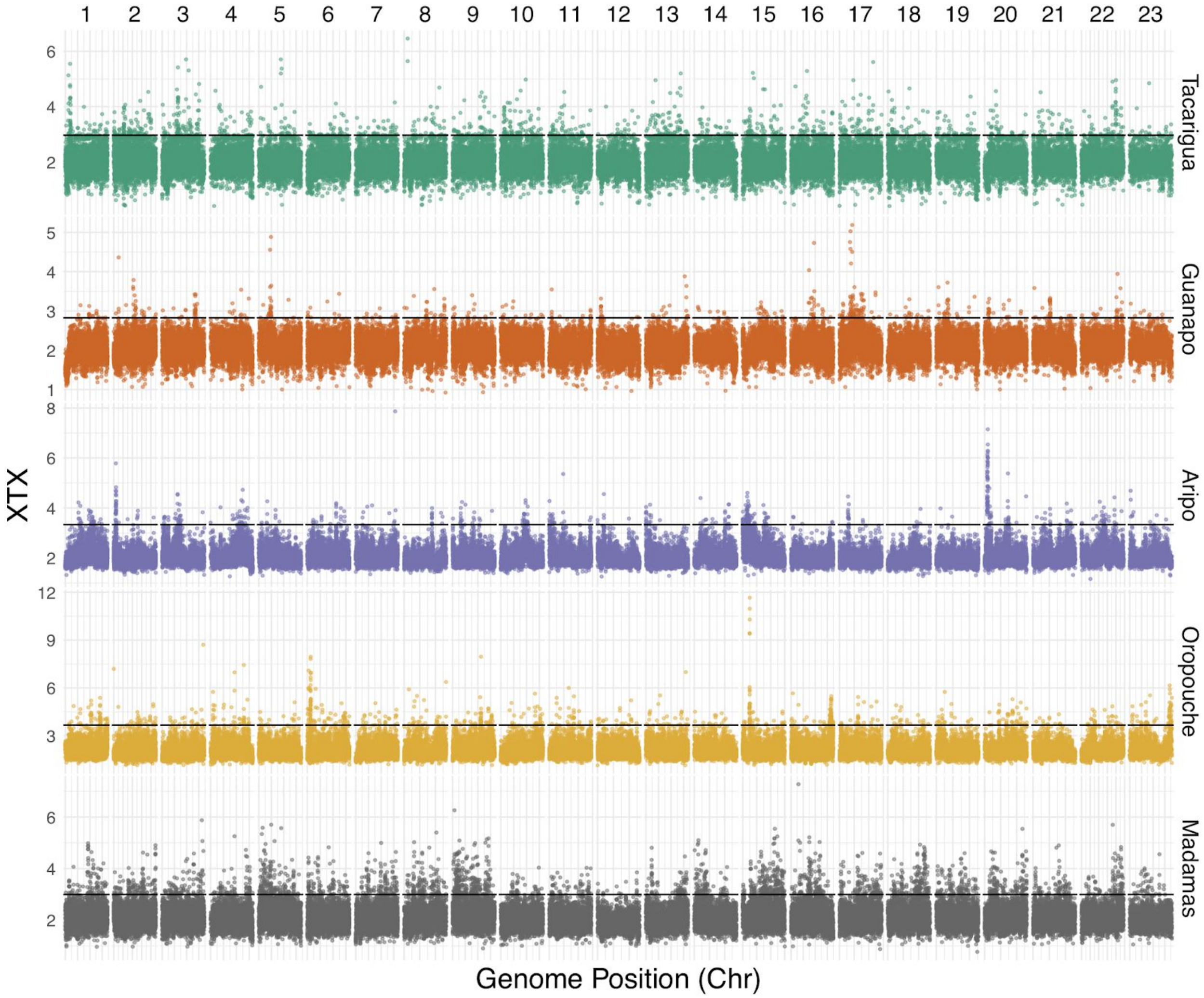


**Figure S5:** Genome-wide XtX results for 10kb windows. Panels represent the 23 chromosomes in the guppy genome. Each row represents the XtX score (a Bayesian analogue of $F_{ST}$, describing relative genetic differentiation) for 10kb windows between HP and LP populations in a different river. Chr20 has been updated to include the unplaced scaffold 000094F. The horizontal line in each row denotes river-specific outlier cutoffs, calculated according to neutral simulations of XtX within each river.

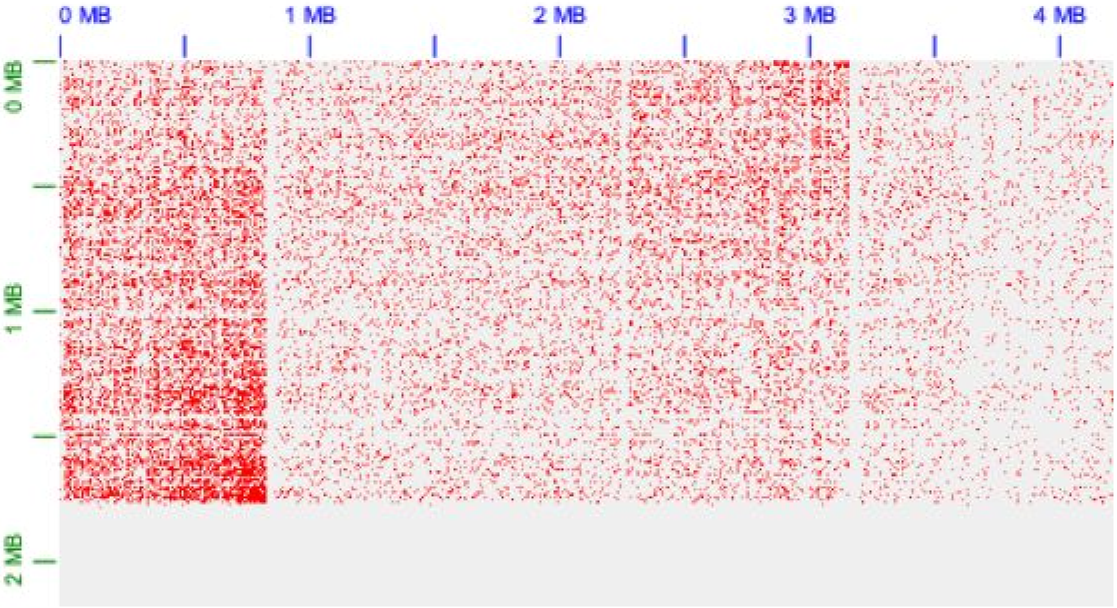


**Figure S6:** HiC contact information used to place scaffold 000094F_0. Chromosome 20 co-ordinates are displayed along the upper axis, and scaffold 000094F_0 co-ordinates along the vertical axis. Scaffold 000094F_0 (total length = 1797025 bp) was reversed and inserted at position 836423 on chromosome 20. The preceding region on chromosome 20 (836423 - 3164071) was also inverted to reflect higher contact between the start of 000094F_0 and the chr20 region around 3164071.

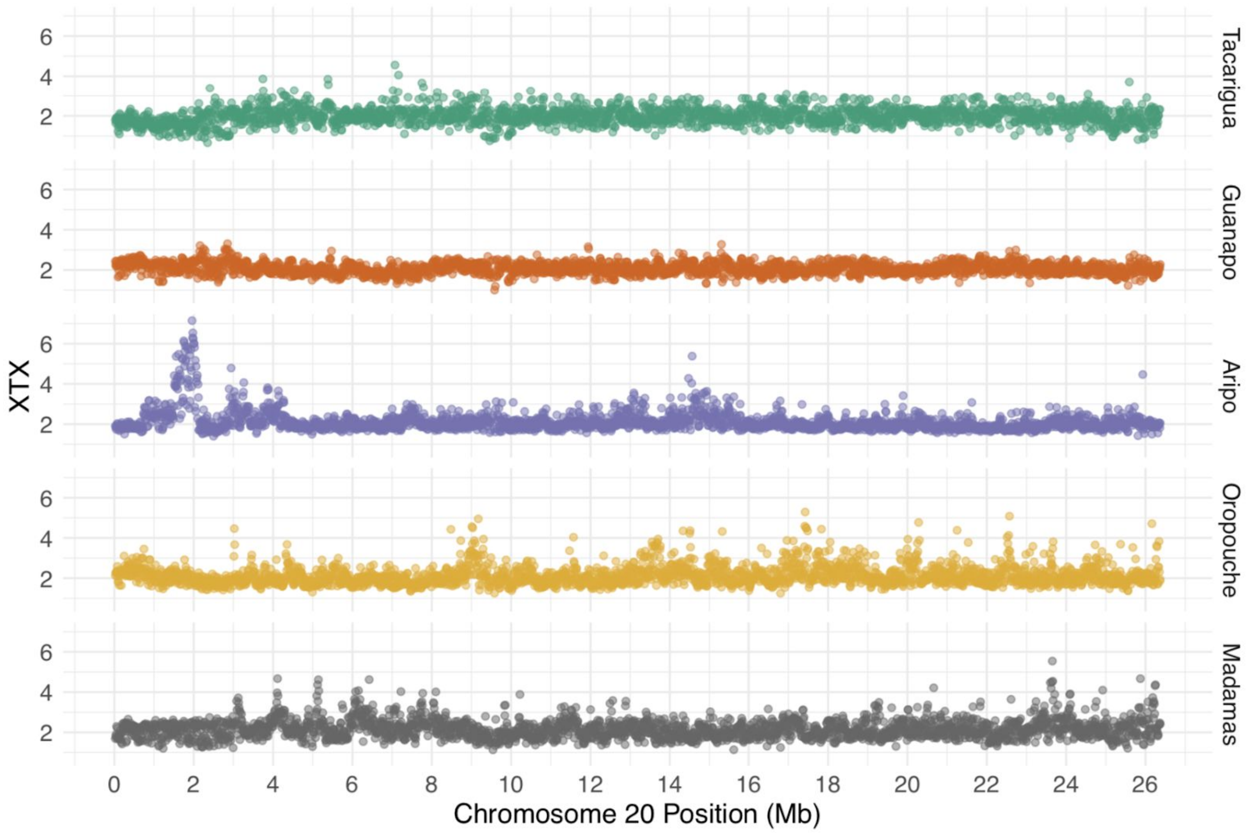


**Figure S7:** XtX results for 10kb windows along chromosome 20 for each river. Each row represents the XtX score (a Bayesian analogue of $F_{ST}$, describing relative genetic differentiation) for 10kb windows between HP and LP populations in a different river. Chr20 has been updated to include the unplaced scaffold 000094F. This figure highlights the location of a peak of strong HP-LP differentiation int the Aripo river.

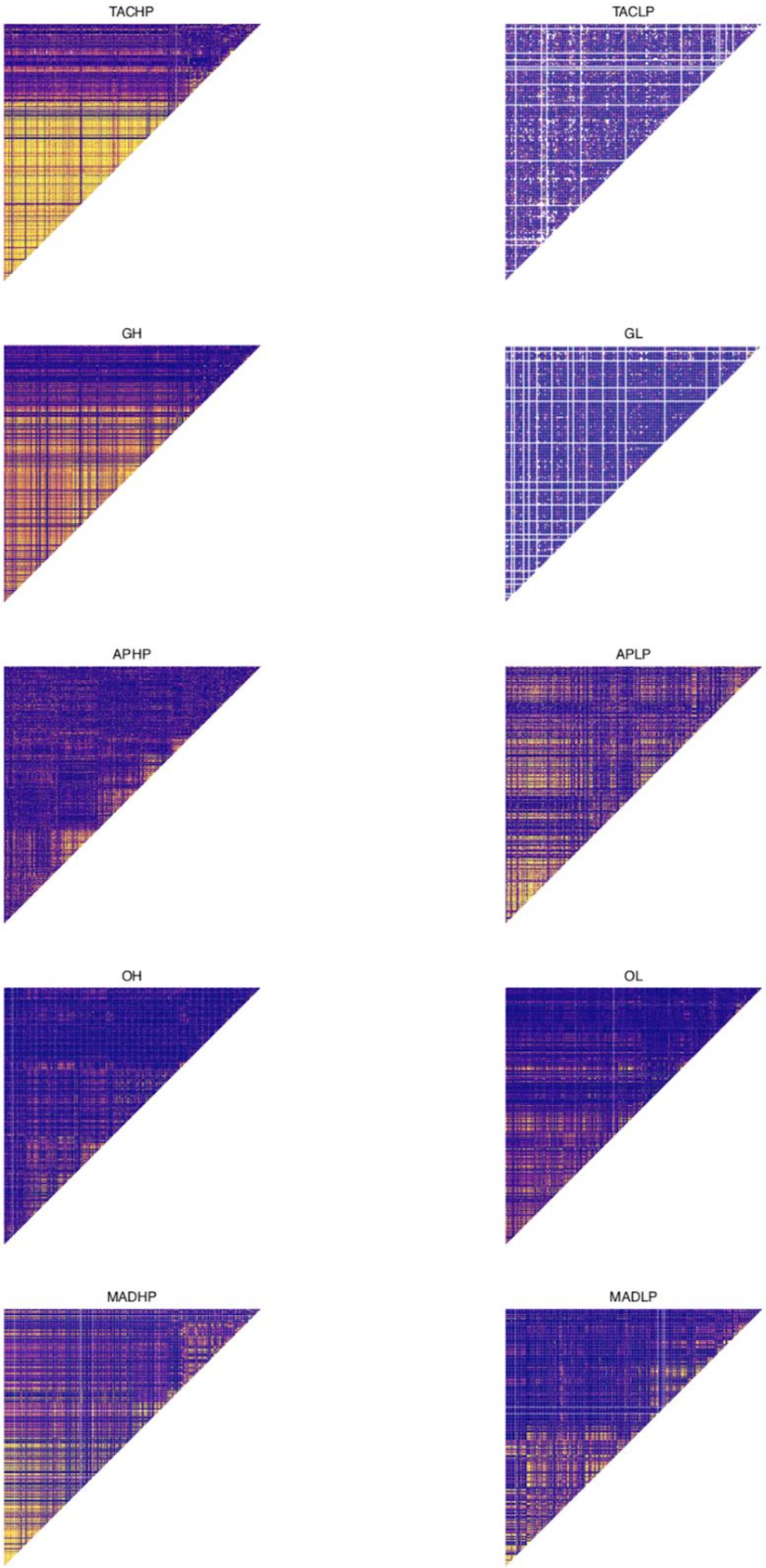


**Figure S8:** Linkage plots across the first 4 Mb of chromosome 20 (following merging of chromosome 20 and scaffold 000094F_0).

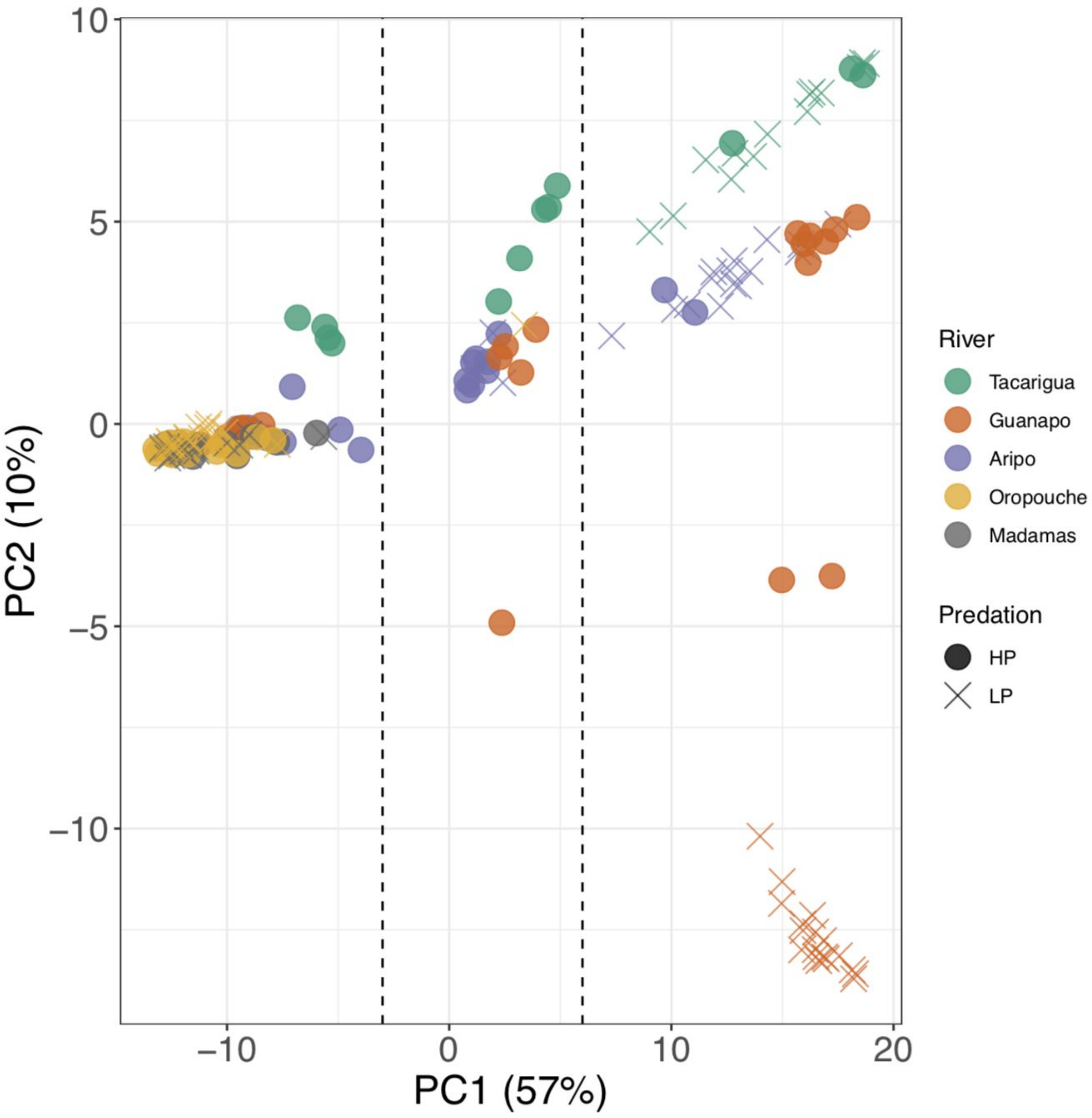


**Figure S9:** PCA analysis across the CL-AP region for all individuals, highlighting three clusters corresponding to homozygotes (REF and CL), and heterozygotes. Dashed lines denote cut-offs used to define haplogroups.

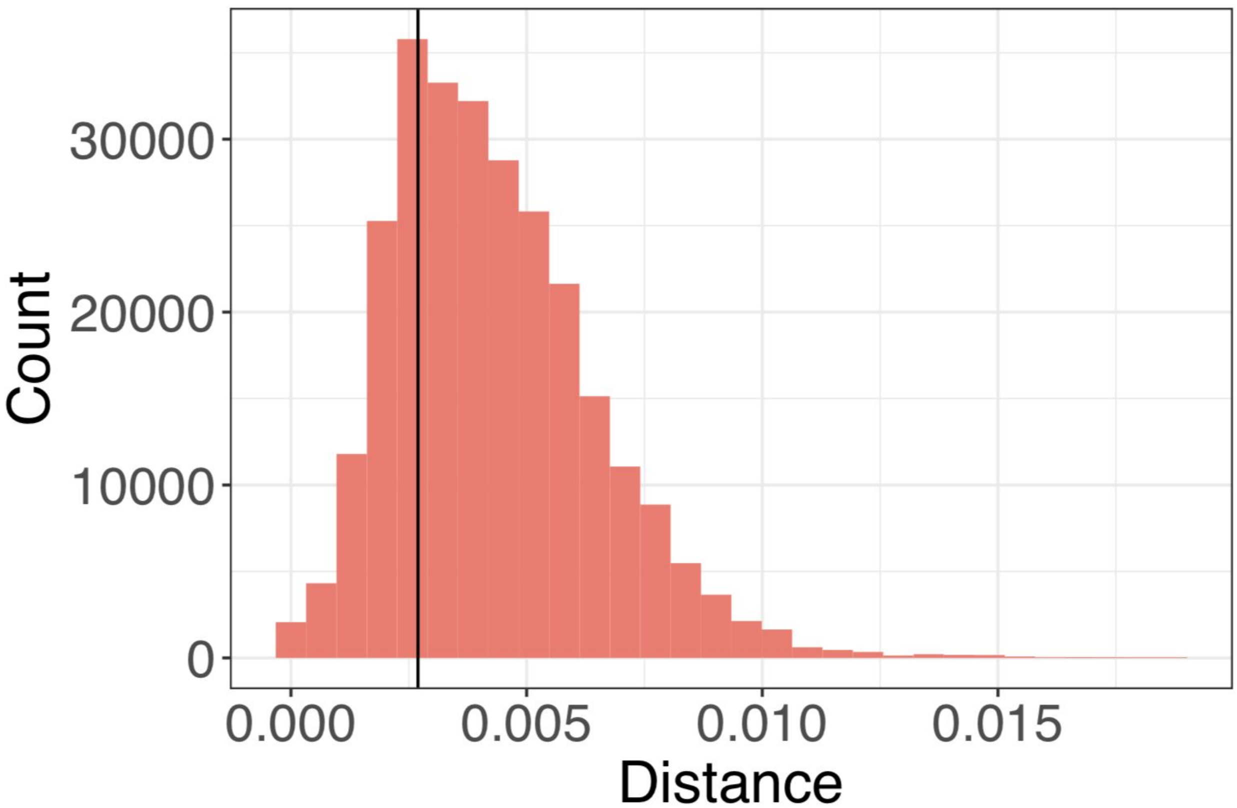


**FIgure S10**: Analysis of branch lengths between Oropouche/Madamas individuals and Caroni individuals (Aripo, Guanapo, Tacarigua) homozygous for the CL haplotype at the CL-AP region. The black line denotes the mean branch distance between haplogroups at the CL-AP region (mean = 2.69e-3). The histogram is comprised of distances estimated between these haplogroups at 50 randomly selected 100kb regions from across the genome. All trees were calculated using RAxML-NG and bootstrapped to a confidence score of <3%.

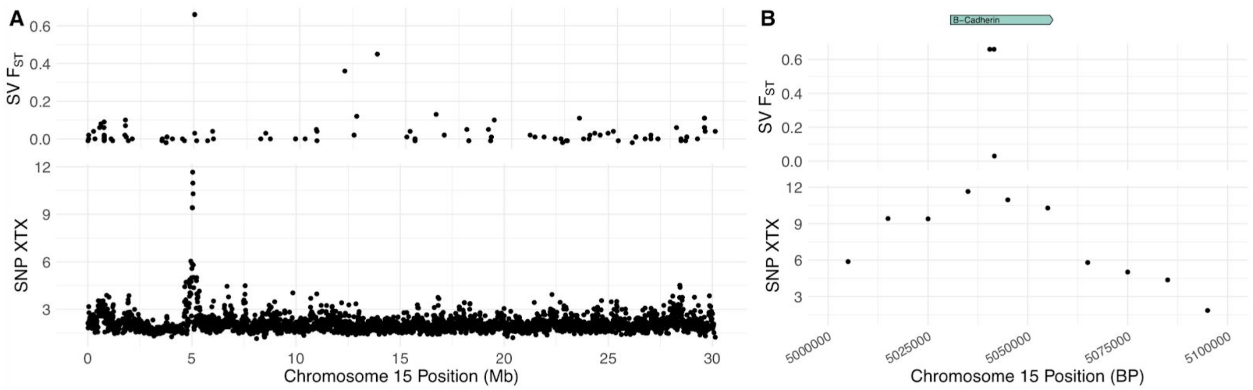


**Figure S11:** Structural variant (SV) $F_{ST}$ and SNP XTX along chromosome 15 in the Oropouche river between HP and LP populations, highlighting concordance between SV and SNP peaks at ~5 Mb **(A).** These peaks corresponded with the *B-cadherin* gene **(B)** in this region. The SV points at this peak correspond with the breakpoints of a 1,097 bp deletion detected using the software *smoove*.

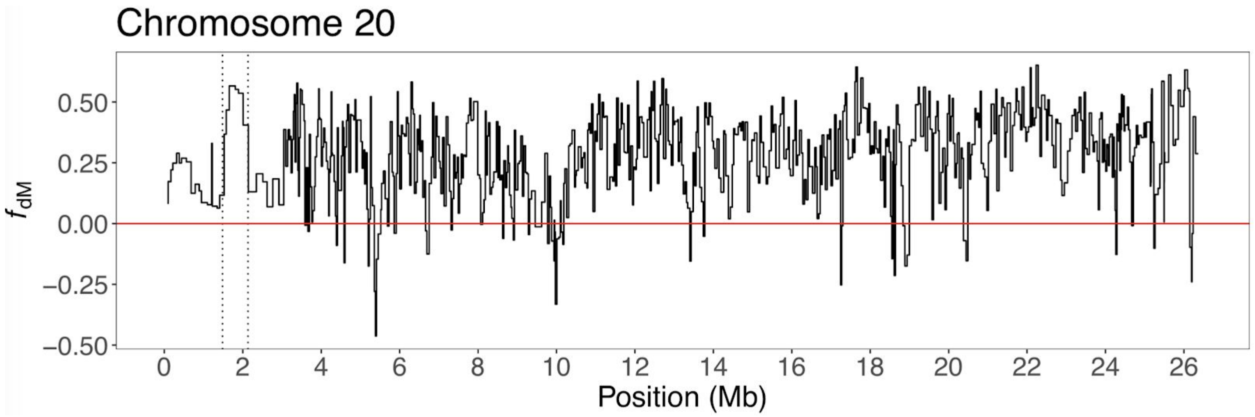


**Figure S12:** Introgression statistics highlighting suspected introgression at the CL-AP region (dotted lines) between OHP and APHP. Red line denotes $f_{dM}$ = 0. Where $f_{dM} < 0$, populations GH and OHP exhibit more shared sites than expected, where $f_{dM} > 0$, populations APHP and OHP exhibit more shared sites than expected. Each plotted point represents a non-overlapping window of 100 SNPs.

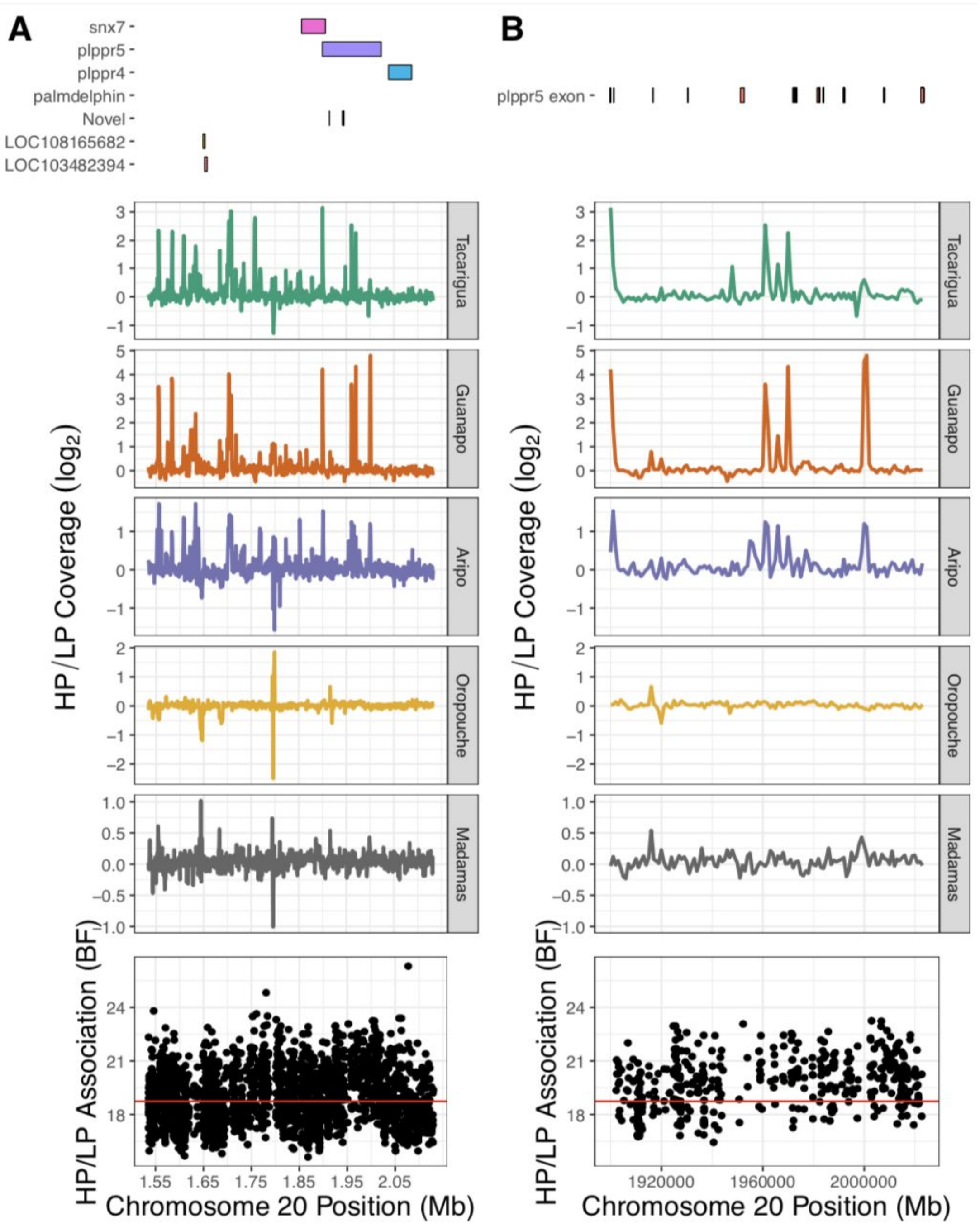


**Figure S13:** Summary of CL-AP region (**A**) and *plppr5* gene region (**B**) according to differences in coverage between HP/LP populations and HP/LP association scores per SNP (BF). Of particular note are a peak in HP/LP coverage ratio in Tacarigua, Guanapo and Aripo at ~1.9 Mb (overlapping the *plppr5* gene), and the SNP with the highest genome-wide HP/LP association score at ~2.06 Mb (overlapping the *plppr4* gene). The peak in HP/LP coverage overlapped with the last exon of *plppr5*, was driven by reduced coverage in LP populations, and was thus confirmed as a ~1kb deletion in the CL haplotype in all Caroni LP populations by visualising bams in igv.
